# MAGIC: Mosaic Analysis by gRNA-Induced Crossing-over

**DOI:** 10.1101/2020.06.26.174045

**Authors:** Sarah E. Allen, Gabriel T. Koreman, Ankita Sarkar, Bei Wang, Mariana F. Wolfner, Chun Han

**Affiliations:** Department of Molecular Biology and Genetics, Cornell University, Ithaca, NY 14853, USA; Weill Institute for Cell and Molecular Biology, Cornell University, Ithaca, NY 14853, USA; These authors contributed equally to this work, listed alphabetically

**Keywords:** *MAGIC*, *mosaic analysis*, CRISPR/Cas9, gRNA, clonal analysis, germline, imaginal disc, da neurons, *Drosophila dominant female sterility*

## Abstract

Mosaic animals have provided the platform for many fundamental discoveries in developmental biology, cell biology, and other fields. Techniques to produce mosaic animals by mitotic recombination have been extensively developed in *Drosophila melanogaster* but are less common for other laboratory organisms. Here, we report <u>m</u>osaic <u>a</u>nalysis by <u>g</u>RNA-<u>i</u>nduced <u>c</u>rossing-over (MAGIC), a new technique for generating mosaic animals based on DNA double-strand breaks produced by CRISPR/Cas9. MAGIC efficiently produces mosaic clones in both somatic tissues and the germline of *Drosophila*. Further, by developing a MAGIC toolkit for one chromosome arm, we demonstrate the method’s application in characterizing gene function in neural development and in generating fluorescently marked clones in wild-derived *Drosophila* strains. Eliminating the need to introduce recombinase-recognition sites in the genome, this simple and versatile system simplifies mosaic analysis in *Drosophila* and can be applied in any organism that is compatible with CRISPR/Cas9.

## Introduction

Mosaic animals contain genetically distinct populations of cells that have arisen from one zygote. Mosaic animals have historically played important roles in the study of pleiotropic genes, developmental timing, cell lineage, neural wiring, and other complex biological processes. Given its genetic tractability, *Drosophila* has been a major system for generating and studying such mosaics [1], which have led to important discoveries such as developmental compartments [2], cell autonomy [3], and maternal effects of zygotic lethal genes [4]. Mosaic (also called clonal) analysis is currently used to study tumor suppressors [5], signaling pathways [6], sleep-wake behaviors [7], cell fates [8], and neuronal lineages [9], among other biological processes.

The earliest mosaic analyses relied on spontaneous mitotic recombination [10], rare events in which a DNA double-strand break (DSB) during the G_2_ phase of the cell cycle is repaired by homologous recombination, resulting in the reciprocal exchange of chromosomal arms between homologous chromosomes distal to the site of the DNA crossover (reviewed in Griffin et al., 2014). Ionizing radiation, such as X-rays [12], cause DSBs and thus were later used in *Drosophila* to increase the baseline level of mitotic recombination [13]. However, ionizing radiation breaks genomic DNA at random locations and is associated with a high degree of lethality.

To overcome these limitations, the yeast Flippase (Flp)/FRT system was introduced into *Drosophila* to mediate site-specific recombination at FRT sites [14,15], enabling the development of an ever-expanding toolbox with enhanced power and flexibility for clonal analysis [15–19]. This system requires that both homologous chromosomes carry FRT sites at the same position proximal (relative to the centromere) to the gene of interest, and an independent marker on one of the homologs to allow visualization of the genetically distinct clones [15]. For clonal analysis in the *Drosophila* germline, a dominant female sterility (DFS) *ovo*^*D1*^ transgene was combined with Flp/FRT methods, allowing production of, and selection for, germline clones homozygous for a mutation of interest in a heterozygous mother [20,21]. In this “Flp-DFS” technique, egg production from *ovo*^*D1*^-containing heterozygous and homozygous germline cells is blocked, resulting in progeny derived exclusively from germline clones lacking *ovo*^*D1*^ that were generated by mitotic recombination [22]. Clonal analysis based on somatic recombination has also been achieved in mice using the Cre-LoxP system and the reconstitution of fluorescent protein genes as markers [23,24]. Despite these successes, site-specific recombination systems have not been widely used for mitotic recombination in model animals beyond *Drosophila melanogaster* due to the challenging task of introducing recombinase-recognition sites into centromere-proximal regions for every chromosome.

Given the power of mosaic animals in biological research, it would be useful to have a more general approach for inducing interhomolog mitotic recombination in any organism, circumventing the challenges just mentioned. The CRISPR/Cas9 system has great potential for extending clonal analysis, because it can create targeted DSBs in the genomic DNA of a wide array of organisms [25]. This binary system requires only the Cas9 endonuclease and a guide RNA (gRNA) that specifies the DNA target site [26], both of which can be introduced into the cell independently of the location of the target site. CRISPR/Cas9-induced DSBs can be repaired either by non-homologous end joining (NHEJ) or homology-directed repair (HDR). So far, most CRISPR/Cas9 applications in animals have been focused on NHEJ-mediated mutagenesis and HDR-mediated gene replacement [27]. Recently, several studies demonstrated that CRISPR/Cas9-induced DSBs can also induce targeted mitotic recombination in yeast and in the germlines of *Drosophila*, houseflies, and tomatoes [28–31], suggesting the possibility of exploiting this property of CRISPR/Cas9 for mosaic analysis. Here, we report <u>m</u>osaic <u>a</u>nalysis by <u>g</u>RNA-<u>i</u>nduced <u>c</u>rossing-over (MAGIC), a novel technique for mosaic analysis based on CRISPR/Cas9. This method can be used to generate mosaic clones in both the *Drosophila* soma and germline. Based on this method, we built a convenient toolkit to generate and label mosaic clones for genes located on chromosome arm 2L. We demonstrate the success of our toolkit for clonal analysis in the soma and the germline and show its applications in analyzing gene functions in neuronal dendrite development. Lastly, we also demonstrate that MAGIC can be used successfully with unmarked wild-derived strains, indicating that this method can be extended to organisms beyond *Drosophila*.

## Results

### Rationale for MAGIC

MAGIC relies on the action of gRNA/Cas9 in a proliferating cell during G_2_ phase to generate a DSB at a specific position on one chromatid of a homologous pair (Figure 1A). The DSB can induce a crossover between this chromatid and a chromatid from the homologous chromosome, resulting in exchange of chromosome segments between the two chromatids at the location of the DSB. During the subsequent mitotic segregation of chromosomes, a 50% chance exists for identical distal chromosome segments to sort into the same daughter cells, generating “twin spots”, which contain two genetically distinct populations of cells homozygous for the chromosome segment distal to the exchange.

**Figure 1.**
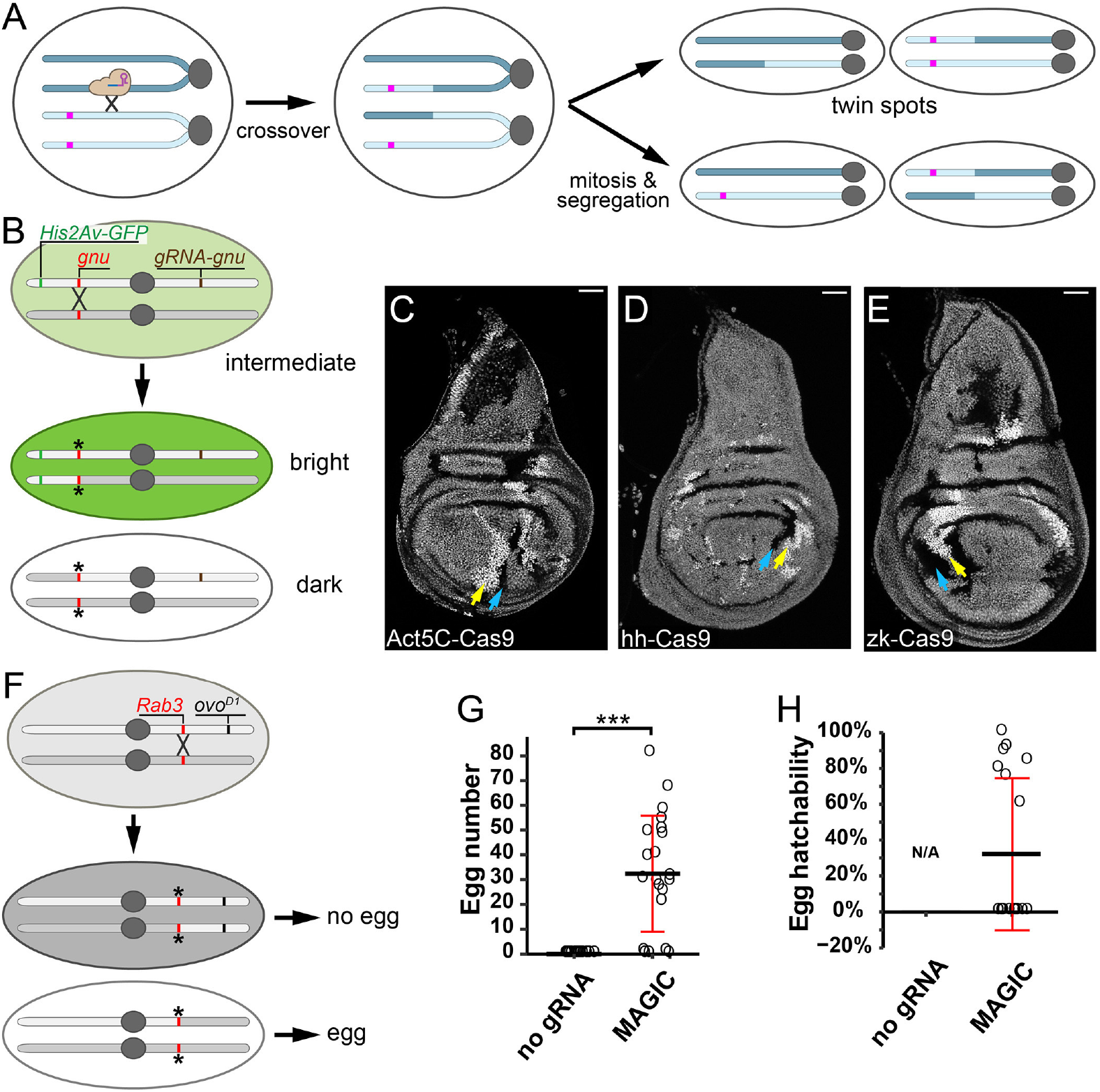
CRISPR-induced crossover generates somatic and germline clones in *Drosophila*. (A) A diagram illustrating how a CRISPR/Cas9-induced DSB leads to generation of homozygous clones in a heterozygous organism. The magenta bar indicates an allele that becomes homozygous in the twin spots. (B) Strategy for generating mosaic clones using the *His2Av-GFP* marker and a crossover at the *gnu* locus. (C-E) Mosaic clones in wing discs visualized by levels of *His2Av-GFP* expression, as described in the text. A pair of arrows indicate *His2Av-GFP*^+/+^ (yellow) and *His2Av-GFP*^−/−^(blue) cells in each panel. Clones were induced by *Act5C-Cas9* (**C**), *hh-Cas9* (**D**) and *zk-Cas9* (**E**). Scale bars, 50 μm. (F) Strategy for generating and detecting clones in the female germline using *ovo*^*D1*^. (G) Number of eggs produced by females carrying *nos-Cas9* and *ovo*^*D1*^, with (MAGIC) or without (no gRNA) *gRNA-Rab3*. ***p<0.001, Student’s t test. n=number females: no gRNA (n=18); MAGIC (n=18). (H) Hatchability of eggs produced females carrying *nos-Cas9* and *ovo*^*D1*^, with (MAGIC) or without (no gRNA) *gRNA-Rab3*. n=number females: no gRNA (N/A); MAGIC (n=18). For all quantifications, black bar, mean; red bar, SD. Asterisks in (B) and (F) indicate mutated gRNA target sites.

### *Using CRISPR-induced crossover to generate clones in the* Drosophila *soma and germline*

For our initial tests of the ability of MAGIC to generate mosaic clones in somatic tissues, we used ubiquitously expressed gRNAs to induce DSBs at the *gnu* locus and a ubiquitous fluorescent marker, *His2Av-GFP* [32], to trace clones (Figure 1B). Both *His2Av-GFP* and *gnu* are located on the left arm of chromosome 3 (3L), and *His2Av-GFP* is distal to *gnu*. We chose *gnu* as our gRNA target because we have already made an efficient *gRNA-gnu* line for other purposes (to be published elsewhere); furthermore, this gene is only required maternally for embryonic development [33], so mutations of *gnu* are not expected to affect the viability or growth of somatic cells. We induced clones using three different Cas9 transgenes, each of which is expressed in the developing wing disc under the control of a different enhancer. With all three Cas9s, we observed twin spots consisting of bright *His2Av-GFP* homozygous clones abutting GFP-negative clones in the midst of *His2Av-GFP/+* heterozygous cells (Figures 1C-1E) in every imaginal disc examined, demonstrating the feasibility of MAGIC for generating somatic mosaics.

Given that CRISPR/Cas9 is active in both the soma and the germline of *Drosophila* [34–36], we next tested for MAGIC clone induction in the germline by using the DFS technique and the germline-specific *nos-Cas9* [35] (Figure S1A). We used an *ovo*^*D1*^ transgene located on chromosome arm 2R and induced DSBs at the *Rab3* locus, which is located on the same arm proximal to the location of *ovo*^*D1*^ (Figure 1F). Since *Rab3* is a non-essential gene expressed only in neurons [37], its disruption in the female germline should affect neither egg production nor embryonic development of the progeny. Due to the dominant effect of *ovo*^*D1*^, restoration of egg production can only result from mitotic recombination proximal to *ovo*^*D1*^ (e.g. at *Rab3* in this case), followed by generation of *ovo*^*D1*^ negative clones. As expected, control females that contained *ovo*^*D1*^ and *nos-Cas9*, but not *gRNA-Rab3*, did not produce any eggs. In contrast, most females carrying all three components produced 20-90 eggs each (Figure 1G), many of which hatched into larvae (Figure 1H), suggesting successful mitotic recombination.

The results above together show that, like the Flp/FRT system, MAGIC is an effective approach for generating homozygous clones via mitotic recombination in both *Drosophila* soma and germline, consistent with the high frequency of CRISPR-induced exchange of chromosomal arms previously demonstrated in the *Drosophila* germline [29]

**Figure S1.**
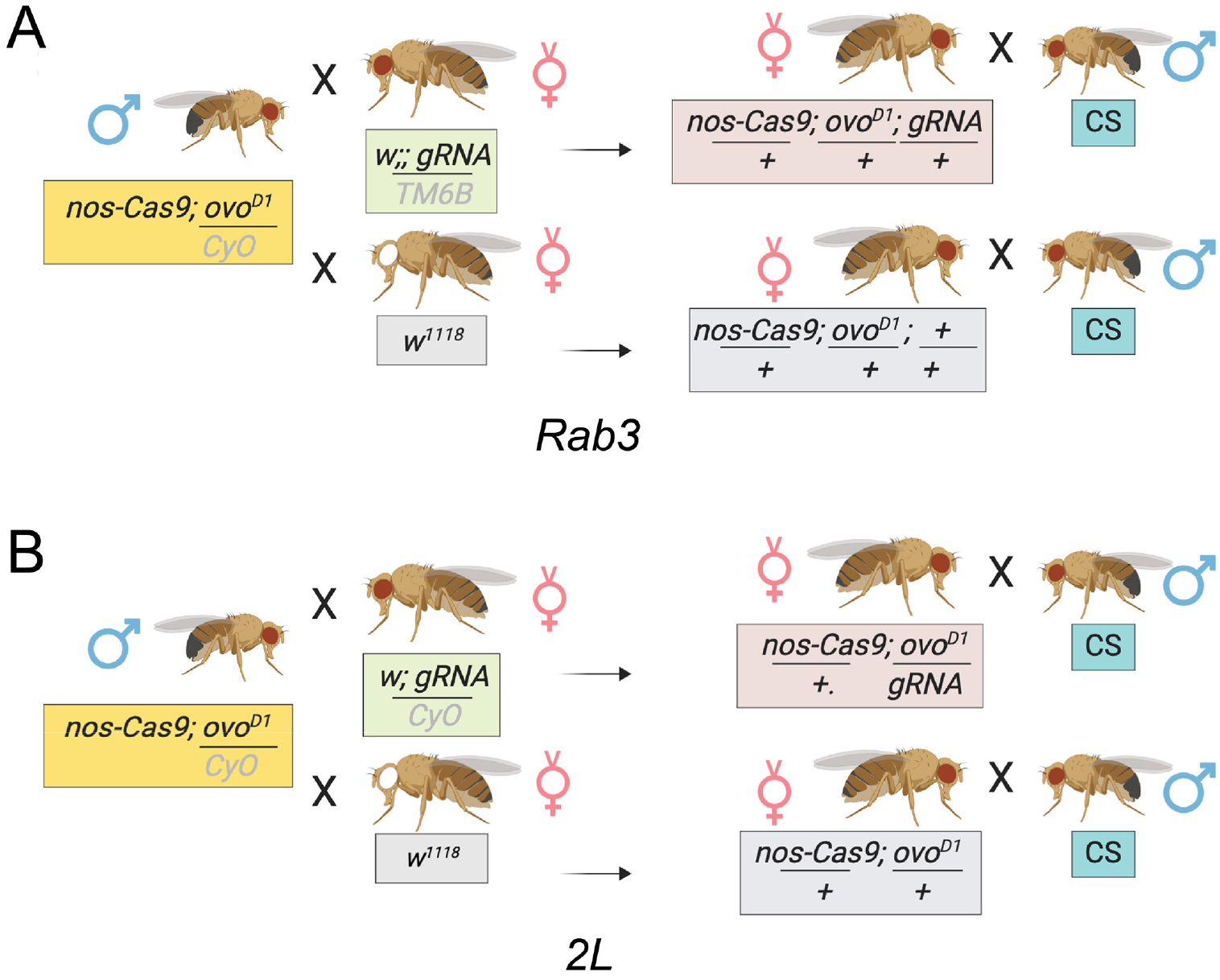
Crossing scheme for germline clone induction. (A) Crossing scheme for germline clone induction using *gRNA-Rab3*, *ovo*^*D1*^(2R), and *nos-Cas9*. (B) Crossing scheme for testing gRNAs for 2L in germline clone induction. The gRNAs used in this test were *gRNA(BFP)* lines.

### A toolkit for generating labeled clones for genes on chromosome arm 2L

Towards making MAGIC a general approach for analyzing *Drosophila* genes, we built a toolkit for genes located on chromosome arm 2L as a proof-of-principle. We designed transgenic constructs that each integrate two features simultaneously: ubiquitously-expressed gRNAs that target a pericentromeric region and a ubiquitously-expressed marker for labeling clones. The constructs were inserted into a distal position of 2L. When used together with an unmodified 2nd chromosome, they label clones homozygous for nearly the entirety of the unmodified arm either negatively or positively. Negative labeling in the nMAGIC option is achieved by expressing a nuclear blue fluorescent protein (nBFP) reporter such that clones homozygous for the unmodified arm lose nBFP expression (Figure 2A). Positive labeling with pMAGIC utilizes a Gal80 marker [17], which suppresses Gal4-driven expression of a fluorescent reporter (Figure 2B). Therefore, only the cells that lose Gal80 transgene will be fluorescently labeled, similarly to mosaic analysis with a repressible cell marker (MARCM) [17].

**Figure 2.**
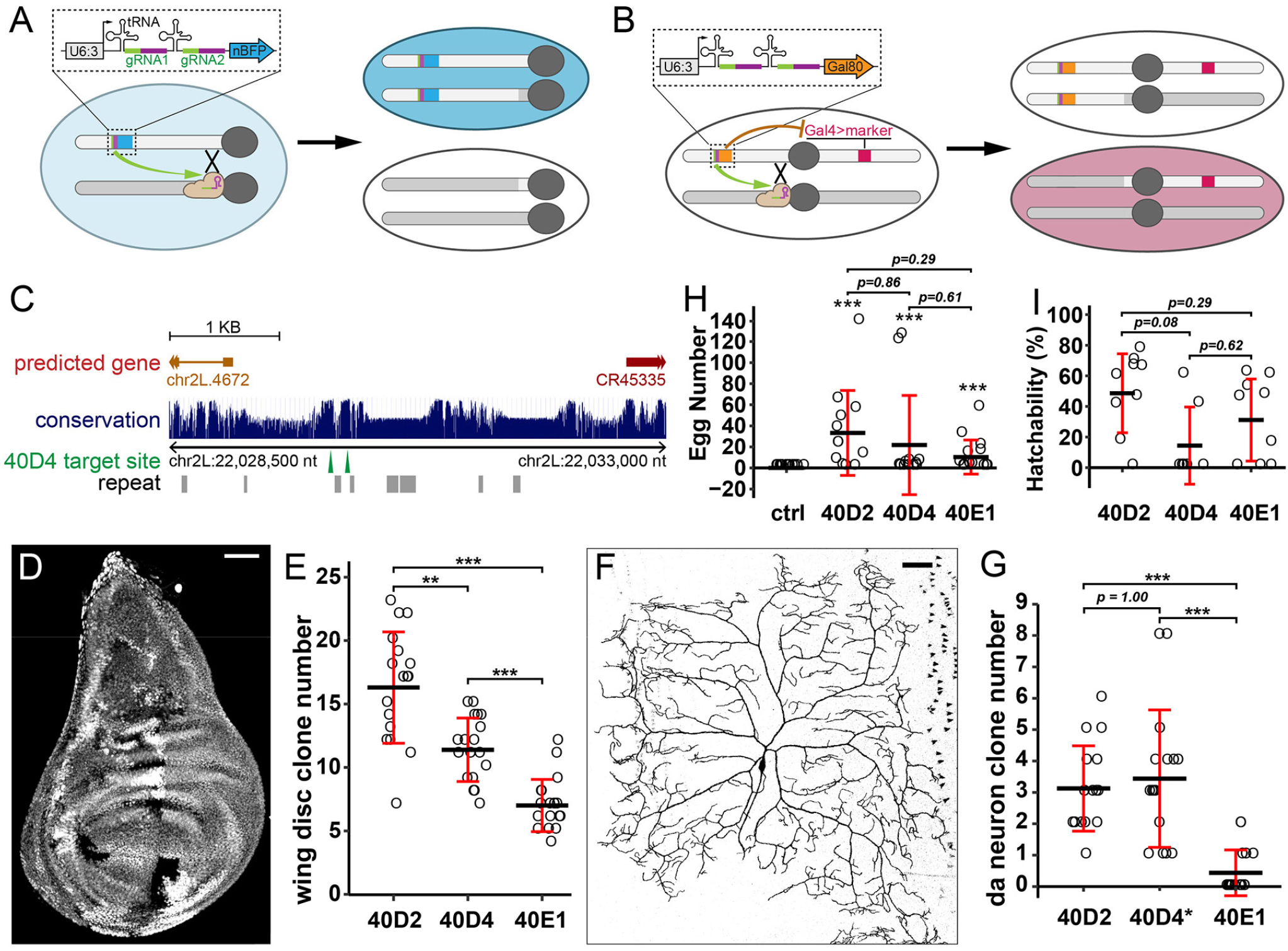
A toolkit for generating labeled clones for genes on chromosome arm 2L. (A) Strategy for generating negatively labeled clones for genes on 2L using nuclear BFP. (B) Strategy for generating positively labeled clones for genes on 2L using Gal80. (C) A map for gRNA-(40D4) target sites derived from the UCSC Genome Browser. The height of the conservation bar corresponds to the level of conservation among 23 *Drosophila* species. (D) A wing imaginal disc showing clones generated by a negative labeling construct (nBFP) paired with *hh-Cas9*. (E) Number of clones in wing discs using three different gRNA(BFP) constructs. n=number of discs: 40D2 (n=17); 40D4 (n=18); 40E1(n=18). Welch’s t-tests, p-values corrected using Bonferroni method; **p≤0.01, ***p≤0.001. (F) A C4da neuron clone generated by a positive labeling construct (Gal80) paired with *SOP-Cas9* and *ppk>CD4-tdTom*. (G) Quantification of C4da clones using three different gRNA(Gal80) constructs. n=number of larvae: 40D2 (n=16); 40D4 (n=16); 40E1 (n=16). Clones were visualized by *ppk>CD4-tdTom.* Welch’s t-tests, p-values corrected using Bonferroni method; ***p≤0.001. The asterisk on 40D4 indicates frequent labeling of C3da clones. (H) Number of eggs produced by using *ovo*^*D1*^(2L), *nos-Cas9*, and three different gRNA(BFP) constructs. The control (ctrl) has no gRNA. n=number of females: ctrl (n=13); 40D2 (n =12); 40D4 (n=12); 40E1 (n=14). Asterisks are from *post hoc* single sample t-tests that compare the estimated marginal mean (EMM) of each gRNA against the control (μ = 0); ***p≤0.001. p-values are from contrasts of EMMs of each gRNA (excluding control). EMMs are based on a negative binomial model. All p-values were corrected using the Tukey method. (I) Hatchability of eggs produced by females carrying *ovo*^*D1*^(2L), *nos-Cas9*, and gRNA(BFP) constructs. n=number female: 40D2 (n=9); 40D4 (n=7); 40E1(n=9). p-values are from contrasts of EMMs of proportions of hatched/non-hatched eggs for each gRNA based on a binomial, mixed-effects model and were corrected using the Tukey method. For all quantifications: black bar, mean; red bar, SD. Scale bars, 50 μm.

To identify appropriate gRNA target sites, we surveyed the pericentromeric sequences of 2L for sequences that met three criteria: (1) being reasonably conserved so that DSBs can be induced in most *Drosophila* strains; (2) not functionally critical and being distant from essential sequences so that indel mutations in nearby regions would not disrupt important biological processes; and (3) unique in the genome, so as to avoid off-target effects. Therefore, for each MAGIC construct, we chose a pair of non-repeat gRNA target sequences in an intergenic region to enhance the chance of DSBs. The two gRNA target sequences are closely-linked to reduce the risk of large deletions (Figure 2C). In addition, we preferentially chose sequences that are conserved among closely related *Drosophila* species (*D. melanogaster*, *D. simulans*, and *D. sechellia*) but not in more distant species. Considering the varying efficiencies of different gRNA target sequences, we selected three pairs of gRNAs targeting three chromosomal locations (40D2, 40D4, and 40E1) and tested their ability to produce clones in wing discs, neurons, and the germline.

Clones were induced in a specific tissue by a Cas9 transgene that is expressed in precursor cells of that tissue. We used *hh-Cas9* [38] for nMAGIC in the wing imaginal disc (Figure 2D), *SOP-Cas9* for pMAGIC in larval class IV dendritic arborization (C4da) sensory neurons (Figure 2F), and *nos-Cas9* for the female germline (Figure S1B). gRNAs targeting 40D2 consistently performed the best in generating clones in wing discs and C4da neurons (Figures 2E and 2G) and appear to be the most efficient in the germline, even though the differences in the germline were not statistically significant (Figures 2H and 2I). Although the overall efficiencies of *gRNA-40D2* and *gRNA-40D4* in inducing clones in da neurons are similar (Figure 2G and Figure S2A), *gRNA-40D4* induced more clones in a different type of neuron (class III). The fact that the three gRNA pairs showed similar trends in their ability to induce clones across different tissues suggests that an efficient gRNA construct for one tissue will likely perform well in other tissues also. These results indicate that we have created an efficient MAGIC toolkit for genes located on chromosome arm 2L. Analogous toolkits could easily be made for any other chromosome arm, using the same methods.

**Figure S2.**
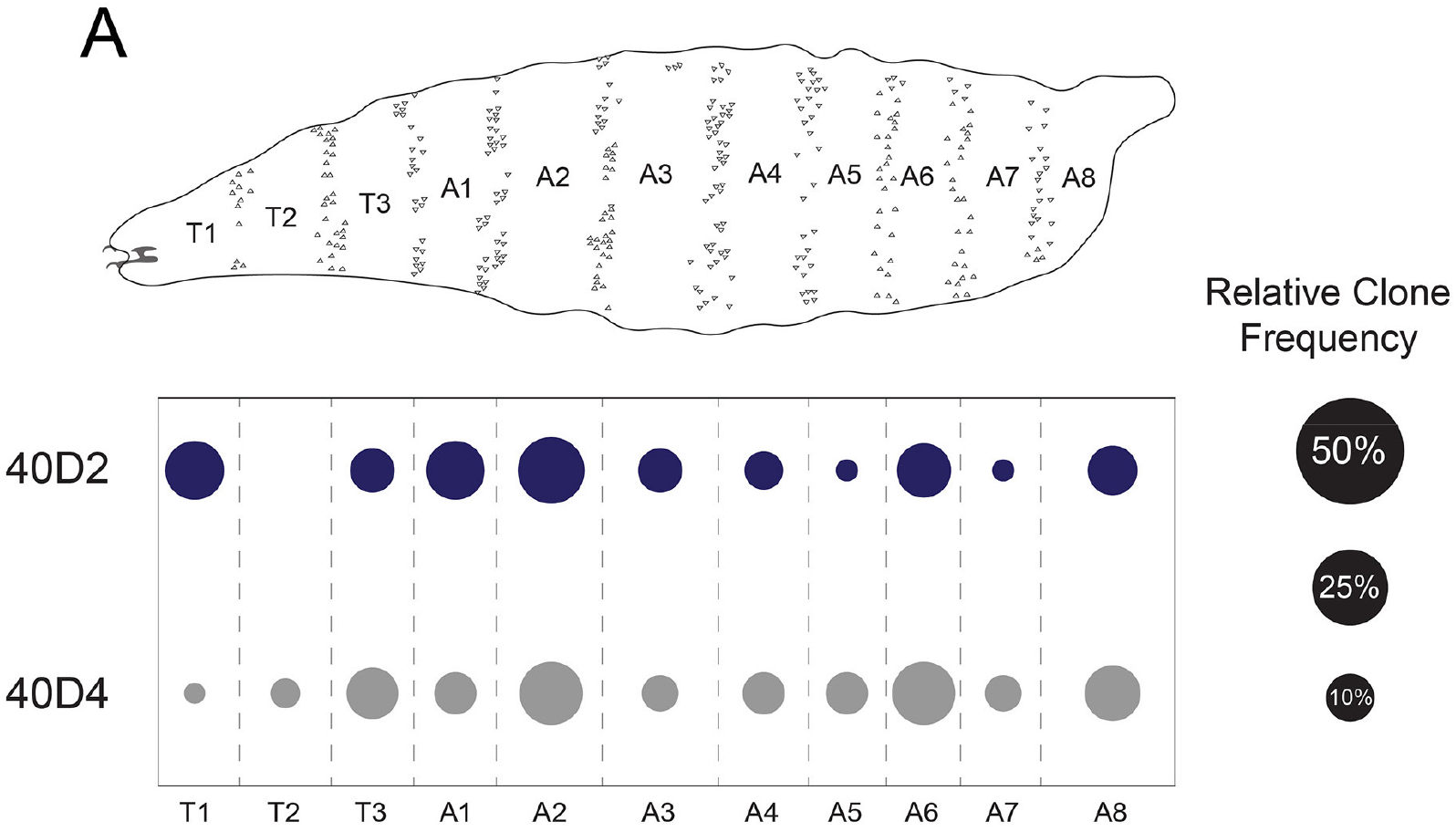
Distribution of da neuron clones using gRNA(Gal80) for 2L. (A) Distribution of da neuron clones in each segment using gRNA(Gal80) for 40D2 and 40D4. n=number of neurons: 40D2 (n=47); 40D4 (n=52).

### Clonal analysis of neuronal dendrite development

To evaluate the utility of our MAGIC toolkit for characterizing gene function at the single-cell level, we combined the pMAGIC line *gRNA-40D2(Gal80)* with mutations on 2L that affect dendrite morphogenesis in C4da neurons by disrupting vesicular trafficking. We first used two genes, *Secretory 5* (*Sec5*) [39] and *Rab5* [40], that have been shown to be required for dendrite growth. We observed dendrite reduction in C4da clones carrying homozygous mutations in these genes (Figures 3A and 3B), recapitulating previously published results using MARCM with the same mutants [39,40]. A third gene, *Syntaxin 5* (*Syx5*), was identified in our unpublished RNAi screens. Clones carrying a null mutation of *Syx5* produced the most dramatic dendrite reduction, with almost all terminal dendrites eliminated (Figure 3C), consistent with the expected role of Syx5 in ER to Golgi vesicle trafficking [41]. Therefore, our MAGIC reagents for 2L can be used to characterize gene functions in single cells with a power analogous to that of MARCM but with a much simpler system.

**Figure 3.**
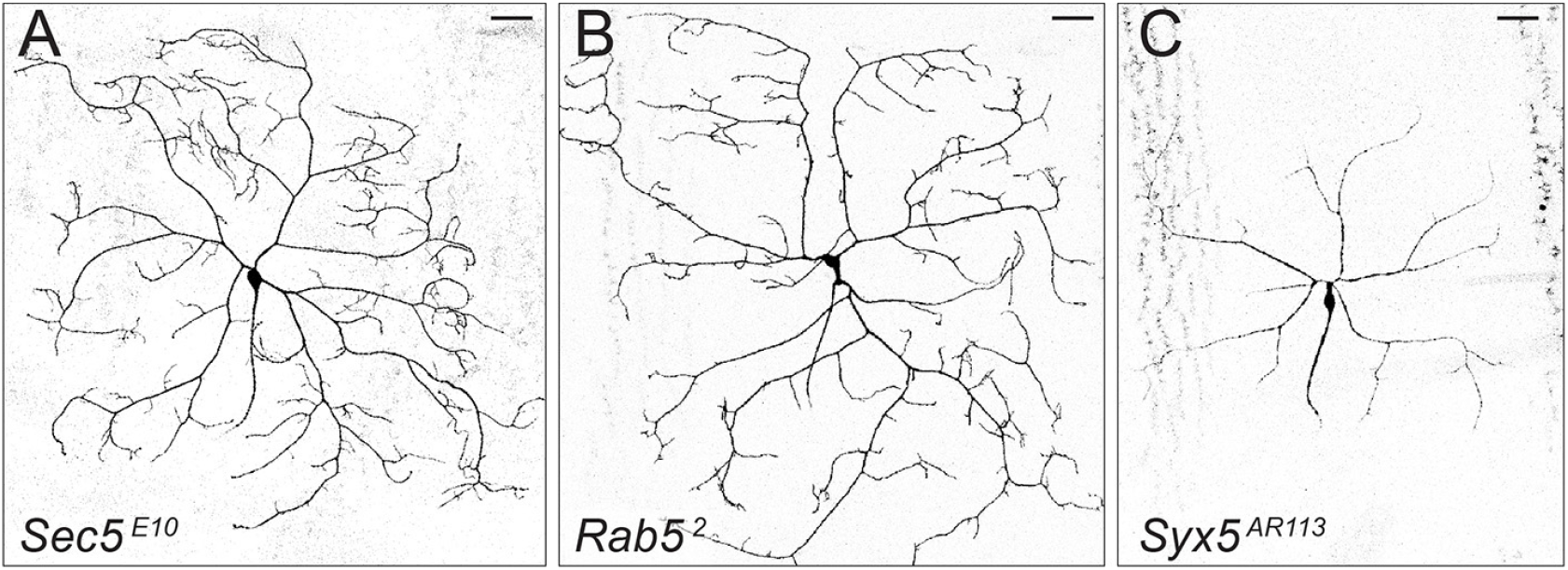
Clonal analysis of example genes in neuronal dendrite development. (A-C) MAGIC clones of C4da neurons using *Sec5*^*E10*^ (A), *Rab5*^*2*^ (B) and *Syx5*^*AR113*^ (C), visualized by *ppk>MApHS*. Scale bars, 50 μm.

### Generation of clones by MAGIC in fly lines with wild-derived genomes

A substantial advantage of MAGIC compared to Flp/FRT-based mitotic recombination systems is that MAGIC does not require prior genetic modification of the chromosome arm to be tested. It therefore has the potential to be applied to fly strains with wild-derived genomes, and even other organisms. To test the applicability of MAGIC to unmarked strains with wild-derived genomes, we crossed *gRNA-40D2(nBFP); hh-Cas9* to five randomly chosen lines from the *Drosophila* Genetic Reference Panel (DGRP) [42], a set of standard strains established from flies captured in the wild. In all cases, we observed efficient clone induction in wing imaginal discs (Figures 4A-4E), demonstrating the potential of MAGIC for clonal analysis of the function of natural alleles residing on wild-derived chromosomes.

**Figure 4.**
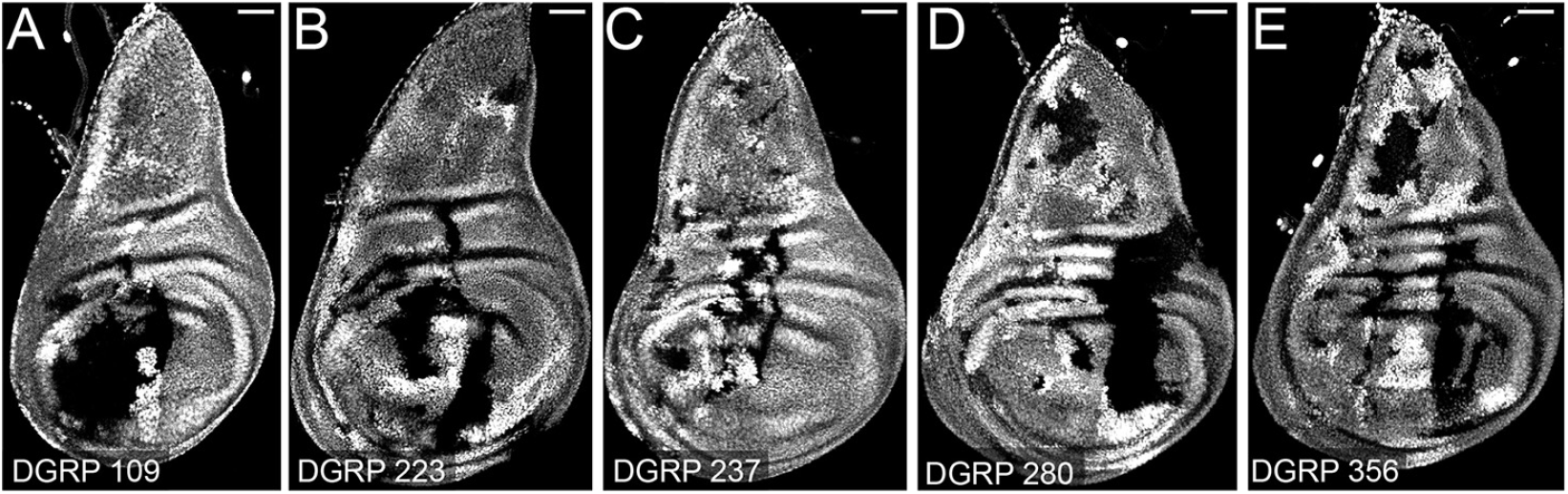
MAGIC generates clones with DGRP lines. (A-E) Clones in wing imaginal discs by pairing *gRNA-40D2(BFP); hh-Cas9* with DGRP line 109 (A), 223 (B), 237 (C), 280 (D) and 356 (E). Scale bars, 50 μm.

## Discussion

We present here a new technique we name MAGIC (<u>m</u>osaic <u>a</u>nalysis by <u>g</u>RNA-<u>i</u>nduced <u>c</u>rossing-over) for clonal analysis based on CRISPR-induced mitotic recombination. We show that MAGIC is capable of producing efficiently mosaic tissues in both the *Drosophila* soma and germline, using gRNAs targeting various chromosomal locations. Integrated gRNA-marker constructs enable both positive- and negative-labeling of homozygous clones. As demonstrated by our 2L toolkit, MAGIC is simple and effective to use; similar MAGIC reagents can be generated easily for all other chromosome arms to allow for genome-wide characterization of gene functions.

Although conventional Flp/FRT-based techniques have been widely and successfully used in *Drosophila* for similar clonal analyses [11], MAGIC has two major advantages. The first is the simplicity of MAGIC assays: This system eliminates the requirement for genetic modification of the test chromosome; furthermore, integrating gRNAs and genetic markers into one transgenic construct further reduces the number of necessary genetic components. Therefore, gRNA-marker transgenes can be combined with existing mutant libraries to perform MAGIC with very little additional effort. The second advantage of MAGIC is that mitotic recombination is not limited by the available FRT insertion sites. While introducing FRT to a specific pericentromeric region is very difficult and historically required labor-intensive genetic screens, CRISPR/Cas9 can induce DNA DSBs and subsequent crossover at specific pericentromeric sequences with ease. Therefore, MAGIC opens doors for clonal analysis of genes that were previously impossible to study using existing FRT sites, such as those on the fourth chromosome [29] and the ones near centromeres. In addition, many *Drosophila* mutations are associated with transgenic constructs containing FRT [43], making analyses complicated when using Flp/FRT-based techniques. In contrast, MAGIC should be compatible with all of these gene disruption lines.

The unique mechanism of MAGIC requires three considerations for successful applications. First, our results suggest that the gRNA target sequence strongly influences the efficiency of clone induction, likely by affecting the frequency of DNA DSBs in premitotic cells. Therefore, for clonal analysis of a specific chromosomal arm, it is beneficial to compare a few candidate gRNA targets and select the most effective one. Second, because perfect DSB repair will recreate the gRNA target site and allow for one more round of Cas9 cutting, most cells that have expressed Cas9 in their lineages are expected to eventually harbor indel mutations that disrupt the gRNA target site, regardless of whether or not the DSBs have led to mitotic recombination. However, this caveat can be mitigated by choosing gRNA sites in non-critical sequences, which can be validated by crossing gRNA lines to a ubiquitous Cas9 or by comparing gRNA-induced control clones to wildtype cells. Lastly, since only DNA DSBs in the G_2_ phase can lead to clone generation, the timing of Cas9 action is expected to be critical for MAGIC. For the cell type in question, an ideal Cas9 should be expressed in the precursor cells, as too early expression can mutate gRNA target sites prematurely and too late expression will lead to unproductive DSBs.

Perhaps the most exciting aspect of MAGIC is its potential for use with wild-derived *Drosophila* strains and in organisms beyond *Drosophila*. DGRP wild-derived strains have played important roles in identification of natural alleles that are associated with certain phenotypic variations [44–47]. However, it has been difficult to investigate the effect of homozygosity for alleles within these strains without being able to use available genetic tools (e.g. Gal4 drivers and fluorescent markers) in *Drosophila*. By combining MAGIC with the DGRP, it is now possible to validate causal effects of specific natural alleles in cellular or developmental processes in a tissue-specific manner. The DGRP can also be used in MAGIC-based genetic screens to identify natural alleles that, when made homozygous, can cause or modify certain phenotypes. Importantly, MAGIC can in theory be utilized in a wide array of organisms that are compatible with CRISPR/Cas9 [48]. In model systems that allow for transgenesis of gRNA-marker constructs, such as mouse, zebrafish, and *Xenopus*, Cas9 can be introduced by injection or virus transduction to further simplify genetic manipulations. Therefore, the flexibility and power of mosaic analysis that are familiar to the *Drosophila* research community are now in reach of researchers who study organisms which have not, or have rarely, been amenable to clonal analysis.

## Materials and Methods

### Fly Stocks and Husbandry

See the Key Resource Table for details of fly stocks used in this study. Broadly, all fly lines were either generated in the Han and Wolfner labs, or obtained from the Bloomington *Drosophila* Stock Center or the *Drosophila* Genetic Reference Panel [42]. All flies were grown on standard yeast-glucose medium, in a 12:12 light/dark cycle, at room temperature (22 ± 1°C, for the egg laying assay) or 25°C (for larval assays) unless otherwise noted. Virgin males and females for mating experiments were aged for 3-5 days. Virgin females were aged on yeasted food for germline clonal analysis.

To test germline clone induction, we combined *nos-Cas9* and *ovo*^*D1*^, and then the gRNA in two sequential crosses in schemes shown in Figure S1.

To visualize clones of C4da neurons, we used *ppk-Gal4 UAS-CD4-tdTom* (Figure 2) and *ppk-Gal4 UAS-MApHS* (Figure 3, only the tdTom channel is shown).

### Molecular Cloning

#### zk-Cas9

The entry vector pENTR221-ZK2 [49] and the destination vector pDEST-APIC-Cas9 (Addgene 121657) were combined in a Gateway LR reaction to generate the expression vector pAPIC2-ZK2-Cas9.

#### MAGIC gRNA-marker vectors

gRNA-marker vectors were constructed similarly to pAC-U63-tgRNA-Rev (Addgene 112811, Poe et al., 2019) but have either a ubi-nBFP (in pAC-U63-tgRNA-nlsBFP) or a ubi-Gal80 (in pAC-U63-tgRNA-Gal80) marker immediately after the U6 3’ flanking sequence. The markers contain a Ubi-p63E promoter, mTagBFP-NLS or Gal80 coding sequence, and His2Av polyA sequence. The Ubi-p63E promoter was amplified from *Ubi-CasExpress* genomic DNA using the oligonucleotides TTAATGCGTATGCATTCTAGTggccatggcttgctgttcttcgcgttc and TTGGATTATTctgcgggcagaaaatagagatgtggaaaattag. mTagBFP-NLS was synthesized as a gBlock DNA fragment (Integrated DNA Technologies, Inc.). Gal80 coding sequence was PCR amplified from pBPGAL80Uw-4 (Addgene 26235) using the oligonucleotides aaaaaaaaatcaaaATGAGCGGTACCGATTACAACAAAAGGAGTAGTGTGAG and GCCGACTGGCTTAGTTAattaattctagaTTAAAGCGAGTAGTGGGAGATGTTG. The His2Av polyA sequence was PCR amplified from pDEST-APLO (Addgene 112805). DNA fragments were assembled together using NEBuilder DNA Assembly (New England Biolabs Inc.).

#### gRNA expression vectors

For *gnu* and *Rab3*, gRNA target sequences were cloned into pAC-U63-tgRNA-Rev as described [38]. For gRNAs targeting 2L, gRNA target sequences were cloned into pAC-U63-tgRNA-nlsBFP and pAC-U63-tgRNA-Gal80 using NEBuilder DNA Assembly. In the gRNA-marker constructs, the tRNA between the first and second gRNAs is a *Drosophila* glutamine tRNA (cagcgcGGTTCCATGGTGTAATGGTTAGCACTCAGGACTCTGAATCCTGCGATCCGAGTTCAAATCTCGGTGGAACCT) instead of a rice glycine tRNA.

Injections were carried out by Rainbow Transgenic Flies (Camarillo, CA 93012 USA) to transform flies through φC31 integrase-mediated integration into attP docker sites. pAPIC2-ZK2-Cas9 and gRNA-marker constructs were integrated into the *attP*^*VK00037*^ site on the second chromosome and expression vectors containing gRNAs targeting *Rab3* or *gnu* were integrated into the *attP*^*VK00027*^ site on the third chromosome. Transgenic insertions were validated by genomic PCR or sequencing.

### Identification of gRNA target sequence

gRNA target sequences for *Rab3* and *gnu* were identified as described previously [38]. Briefly, two gRNA prediction methods were used: sgRNA Scorer 2.0 [50] (https://crispr.med.harvard.edu) and Benchling (www.benchling.com). Candidate target sequences were those that obtained high on-target scores in both algorithms. CasFinder [51] was used to identify and reject any sequences with more than one target site. Two target sequences against coding exons for all splice isoforms were chosen for each targeted gene. gRNA target sequences for 2L were identified by visually scanning through pericentromeric sequences using UCSC Genome Browser (https://genome.ucsc.edu/) following principles described in the Results section. The gRNA target sequences are listed in the table below.

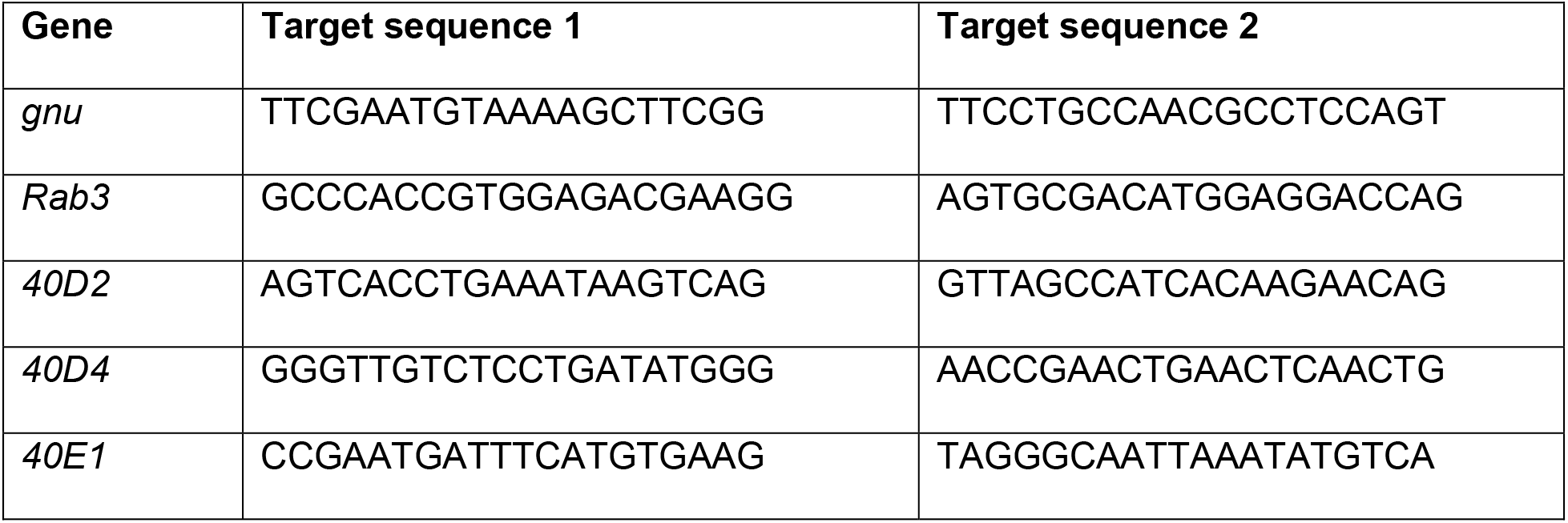

### Live Imaging of neurons

Live imaging was performed as previously described [49]. Briefly, animals were reared at 25°C in density-controlled vials for between 96 and 120 hours after egg-laying (to obtain third to late-third instar larvae). Larvae were mounted in glycerol and their C4da neurons at segments A1-A6 were imaged using a Leica SP8 confocal microscope with a 20x oil objective and a z-step size of 3.5 μm.

### Imaginal disc imaging

Imaginal disc dissections were performed as described previously [52]. Briefly, wandering third instar larvae were dissected in a small petri dish filled with cold PBS. The anterior half of the larva was inverted and the trachea and gut were removed. The sample was then transferred to 4% formaldehyde in PBS and fixed for 15 minutes at room temperature. After washing with PBS, the imaginal discs were placed in SlowFade Diamond Antifade Mountant (Thermo Fisher Scientific) on a glass slide. A coverslip was lightly pressed on top. Imaginal discs were imaged using a Leica SP8 confocal microscope with a 20X oil objective.

### Assays for Germline Clonal Analysis

To monitor mitotic recombination events resulting in germline clone generation, we performed egg-laying and egg hatchability assays as detailed in Hu and Wolfner (2019), with the exception of using Canton-S males in place of ORP2 males as wild-type mates. Hatchability was calculated only for females that laid eggs. Females that laid no eggs were eliminated from hatchability calculations to avoid inflation of false-zero values.

### Image Analysis and Quantification

Counting of wing disc clones was completed manually in Fiji/ImageJ. Counting of neuronal clones was completed manually during the imaging process.

### Statistical Analysis

Statistical analyses were performed in R. Student’s t-test was conducted for egg-laying data using *Rab* and *kni* gRNAs. For egg-laying data using the 2L toolkit, we performed estimated marginal means contrasts between gRNAs and *post hoc* one sample t-tests using a generalized linear model with a negative binomial response. For hatchability data using the 2L toolkit, we performed estimated marginal means contrasts between proportions of hatched/non-hatched eggs for each gRNA using a generalized linear mixed-effects model with a binomial response. For all contrasts, p-values were corrected for multiple comparisons using the Tukey method. For wing disc and neuronal clone data, we performed Welch’s analysis of variance (ANOVA) followed by pairwise *post hoc* Welch’s t-tests. p-values from the multiple *post hoc* Welch’s t-tests were corrected for multiple comparisons using the Bonferroni method.

## Acknowledgments

We thank Bloomington *Drosophila* Stock Center (NIH P40OD018537) and the *Drosophila* Genetic Reference Panel for fly stocks; Cedric Feschotte, Andy Clark, Eric Alani, and Marcus Smolka for advice on gRNA design; Cornell CSCU consultants Stephen Parry and Erika Mudrak for advice on statistics; Michael Goldberg, John Schimenti, Erich Brunner, and Konrad Basler for critical reading and suggestions on the manuscript. This work was supported by a Cornell start-up fund and NIH grants (R01NS099125 and R21OD023824) awarded to C.H., an NIH grant (R01-HD038921) awarded to M.W., and an NIH training grant (T32-GM07273) awarded to the Cornell BMCB graduate program that supported S.A.

---

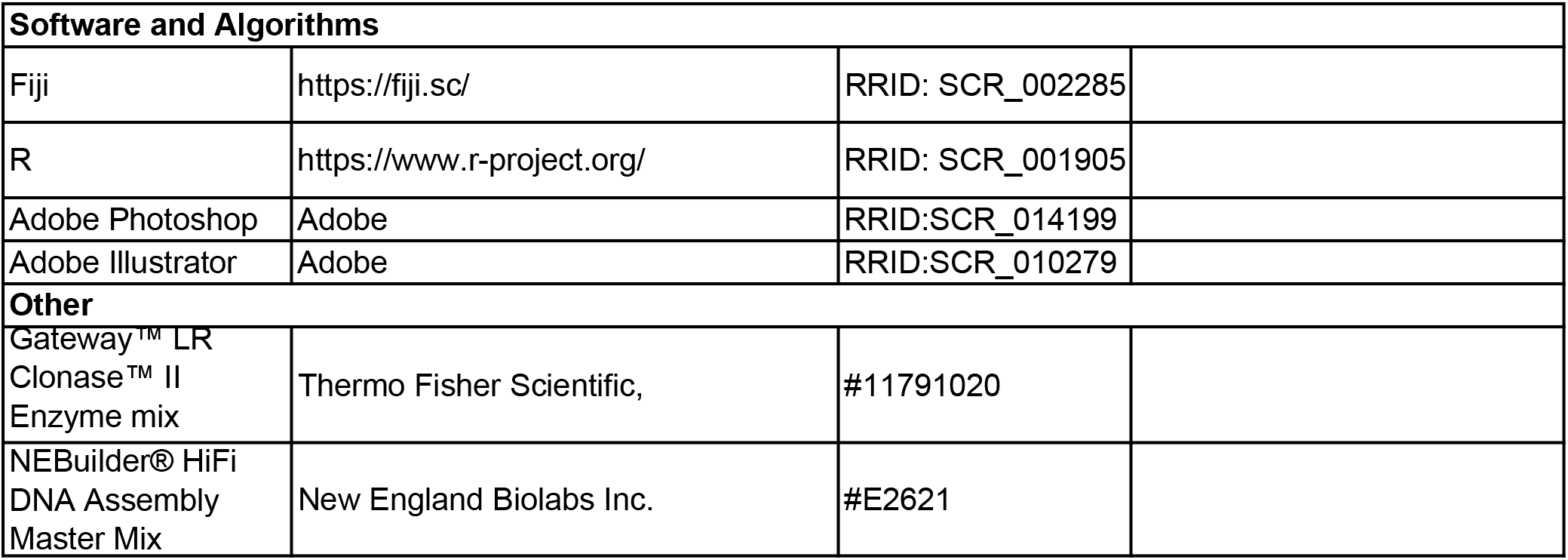
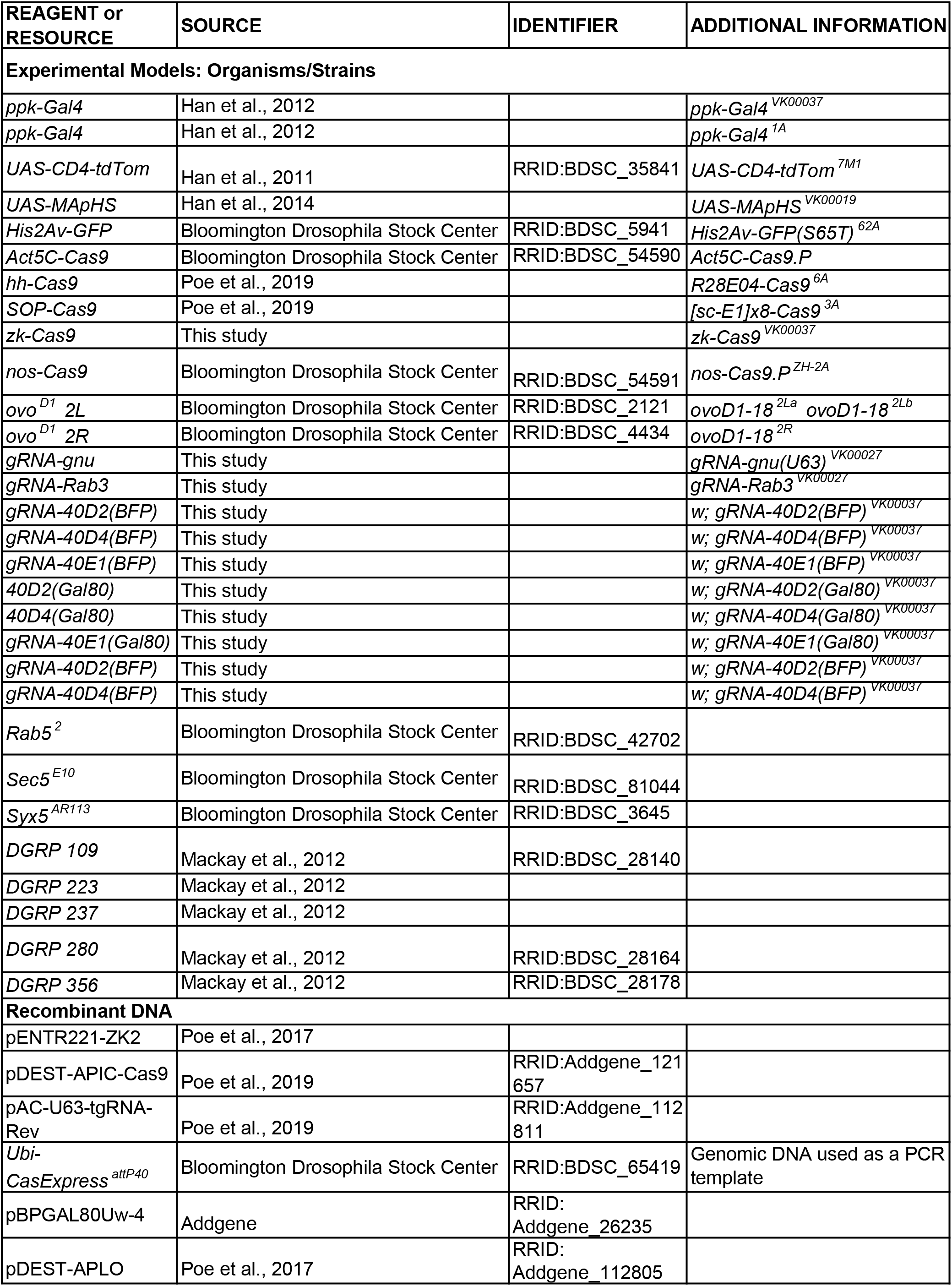
Key Resource Table.

